# Whole genome amplification and sequencing of low cell numbers directly from a bacteria spiked blood model

**DOI:** 10.1101/153965

**Authors:** Catherine Anscombe, Raju.V Misra, Saheer Gharbia

## Abstract

Whilst next generation sequencing is frequently used to whole genome sequence bacteria from cultures, it’s rarely applied directly to clinical samples. Therefore, this study addresses the issue of applying NGS microbial diagnostics directly to blood samples. To demonstrate the potential of direct from blood sequencing a bacteria spiked blood model was developed. Horse blood was spiked with clinical samples of *E. coli* and *S. aureus*, and a process developed to isolate bacterial cells whilst removing the majority of host DNA. One sample of each isolate was then amplified using ϕ29 multiple displacement amplification (MDA) and sequenced. The total processing time, from sample to amplified DNA ready for sequencing was 3.5 hours, significantly faster than the 18-hour overnight culture step which is typically required. Both bacteria showed 100% survival through the processing. The direct from sample sequencing resulted in greater than 92% genome coverage of the pathogens whilst limiting the sequencing of host genome (less than 7% of all reads). Analysis of *de novo* assembled reads allowed accurate genotypic antibiotic resistance prediction. The sample processing is easily applicable to multiple sequencing platforms. Overall this model demonstrates potential to rapidly generate whole genome bacterial data directly from blood.

## Introduction

Bacterial sequencing of clinical isolates has been used in many settings, including for virulence determinants^1–4^, population structure^5,6^ and outbreak investigations ^7–10^. Almost exclusively current applications of sequencing rely either upon culture or targeted amplification imparting a diagnostic bias towards known pathogens and bacteria with known and relatively simple growth requirements. Bacterial whole genome sequencing is most often performed at reference or research centres to study population structures and evolution, with results rarely available in a clinically actionable time. One exception to this is the application of next generation sequencing to the diagnosis of tuberculosis^11,12^, where WGS was shown, in some cases, to be quicker than current routine methods for predicting resistance; however this process still relies on culturing of the bacteria prior to sequencing.

Direct from sample sequencing of pathogens using an untargeted (metagenomic) approach has the potential to lower turnaround times and lower diagnostic bias. Metagenomic sequencing techniques have already been applied to ecological studies ^13–16^, gut microbiomes ^17–19^ and investigation and identification of viruses ^20–23^.

Bacteraemia is a major global cause of morbidity and mortality, with a large range of aetiologies and is a particular problem in healthcare settings^24,25^. The current microbiological diagnostic process is culture based often involving specialised equipment, with time to positivity varying due to aetiology and pathogen load. Although recent advances in MALDI-TOF processing has allowed direct identification of bacteria from positive blood cultures ^26,27,28^, antibiotic sensitivities take a further 18 hours from pure bacterial cultures. The direct application of whole genome sequencing to blood samples would allow rapid pathogen diagnosis, along with simultaneous pathogen typing and genotypic resistance prediction. Furthermore, by applying an unbiased pathogen detection method the diagnostic bias would be lowered.

A two-stage host cell removal was performed, firstly red blood cells were removed using HetaSep^®^, previously used to isolate nucleated cells in the blood, particularly granulocytes. Secondly a selective white blood cell lysis was undertaken using saponin^29^ in order to release intracellular pathogens and aid in host nucleic acid removal.

To directly sequence pathogens, present at low levels (as few as 1 bacterial cell per ml blood) such as those found in sterile site infections an unbiased and high-fidelity amplification enzyme is needed. Multiple displacement amplification (MDA) using ɸ 29 is an alternative method to PCR for the production of DNA in high enough amounts for sequencing. ɸ 29 MDA has been applied to samples with very low starting DNA amounts from single cells (both prokaryotic and eukaryotic) and provided DNA in levels high enough to perform sequencing ^30,31^. It has been demonstrated that this method is suitable for use with single bacterial cells^30,31^ and more recently malaria parasites directly from blood samples ^32^. The aim of this study was to devise a process to allow same day untargeted bacterial sequencing to be performed on DNA directly isolated from a bacteria spiked blood model.

## Methods

### Isolates

Clinical isolates of *E. coli* and *S. aureus* were collected from the Royal Free Hospital Hampstead, where they had been stored at -80°C after isolation from septic patients. *E. coli* and *S. aureus* are common aetiologies of blood stream infections, representing different cell wall types and cell morphologies. Phenotypic data was produced using the BD phoenix and was retrieved from final hospital reports. Bacteria were cultured on Horse blood agar at 37°C overnight.

### Spiking and isolation process

Spiked samples were set up by adding an estimated ten bacterial cells of the clinical isolate of *S. aureus* or *E. coli* calculated using serial dilutions to 1ml horse blood. The workflow depicted in **Error! Reference source not found.** Figure 1 was then applied, 200 μl HetaSep^®^ was added and the sample vortexed and incubated at 37°C for 10 minutes. 550 μl supernatant was removed and 200 μl 5% saponin was added to a final 2% solution, and incubated at room temperature for 5 minutes. 700μl sterile water was added for a water shock and incubated at room temperature for 30 seconds before salt restoration with the addition of 21μl 5M NaCl. The sample was centrifuged at 4000xg for 5 minutes and the supernatant discarded before addition of 2 μl turbo DNase1 and 5 μl of 10x buffer (Ambion). The sample was vortexed and incubated at 37°C for 15 minutes EDTA was added to a final concentration of 15nM. The sample was then centrifuged for 5 minutes at 4000xg and the supernatant removed and discarded. The pellet was washed in decreasing volumes of PBS, initially 200 μl then 100 μl followed by 20 μl with each stage being centrifuged at 6000xg for three minutes. The final pellet was suspended in 4 μl sterile PBS.

**Figure 1.**
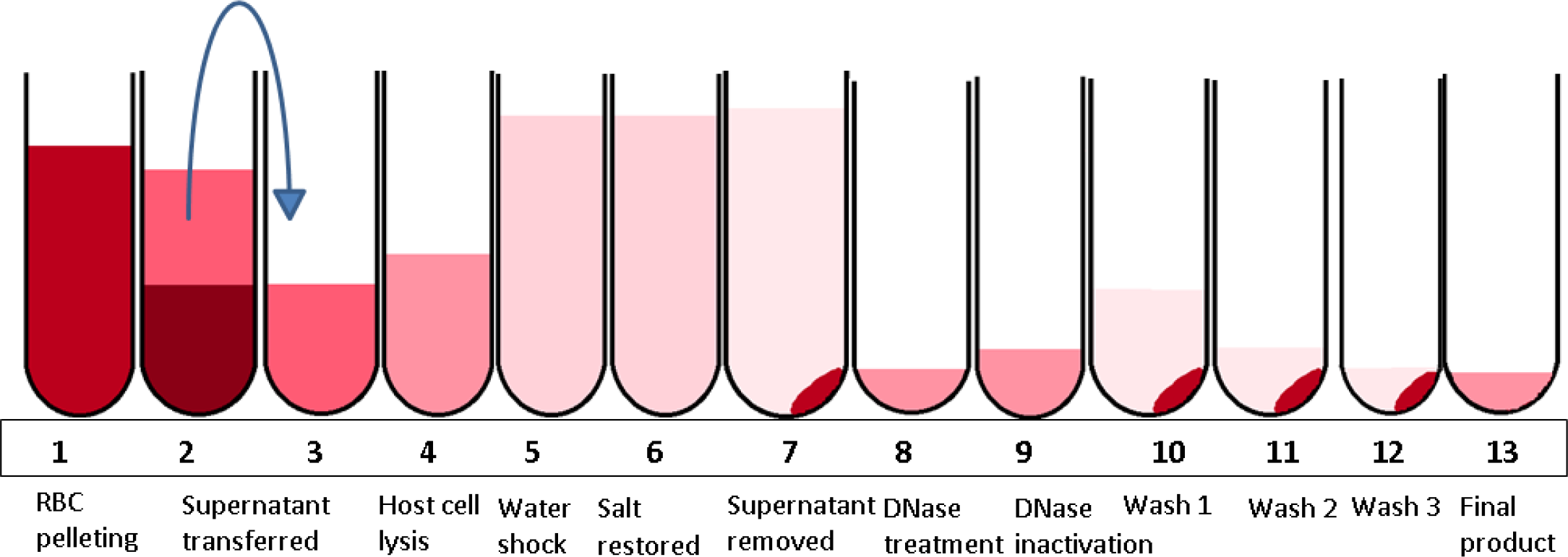
Illustration of the process for bacterial isolation from whole blood using HetaSep and selective lysis with Saponin. Numbers refer to sampling points where bacterial recovery was investigated. Total processing time for bacterial isolation is one hour.

### Bacterial survival

To check for bacterial survival at each stage of processing 13 spiked samples were set up and processed. At the end of each processing stage (Table 1) one sample was cultured and colony counts performed. This was repeated three times for the *E. coli* and *S. aureus* isolate.

**Table 1.** the average number of CFU isolated at each processing stage for bacterial isolation from whole blood using HetaSep and selective lysis with Saponin after workflow improvements. Numbers refer to sampling points where bacterial recovery was investigated as illustrated in **Figure 1**

| Stage |  | *S. aureus* | *E. coli* |
| --- | --- | --- | --- |
| 1 | Spiked cells | 10 | 11 |
| 2 | RBC separation-bottom | 0 | 0 |
| 3 | RBC separation-top | 10 | 9 |
| 4 | Addition of Saponin | 12 | 9 |
| 5 | Water shock | 12 | 12 |
| 6 | Salt restoration | 10 | 10 |
| 7 | Cell pelleting-top | 0 | 0 |
|  | Cell pelleting-bottom | 9 | 9 |
| 8 | DNAse treatment | 11 | 12 |
| 9 | EDTA and PBS | 10 | 12 |
| 10 | Wash 1-top | 0 | 0 |
|  | Wash 1-bottom | 9 | 11 |
| 11 | Wash 2-top | 0 | 0 |
|  | Wash 2-bottom | 10 | 10 |
| 12 | Wash 3-top | 0 | 0 |
|  | Wash 3-bottom | 8 | 9 |
| 13 | Final pellet | 11 | 9 |

### DNA extraction and Amplification

Extraction was performed using alkaline method; briefly cell suspensions were added to 200 mM potassium hydroxide (Qiagen) and 50mM dithiothreitol (Qiagen) and incubated at 65°C for 10 minutes. The reaction was then neutralised using neutralisation buffer (Qiagen), briefly vortexed and placed on ice.

Amplification was performed using ɸ29 MDA (Repli-g Single Cell Kit Qiagen). A master mix was prepared on ice in a total volume of 40 µl, with 29 µl reaction buffer, containing endonuclease resistant hexamer primers and 2 µl (40 U) of ɸ29 polymerase (Qiagen, REPLI-g Single Cell Kit). The extracted DNA was then added to the master mix and the sample was then incubated at 30°C for 2 hours, and the reaction stopped by heating to 65°C for 3 minutes.

### Sequencing

DNA was quantified using Qubit BR kit and 3 µg of ɸ29 MDA DNA was de-branched using S1 nuclease in a 90 µl reaction as follows, 3 µl 10x buffer, 3 µl 0.5M NaCl, 10 µl S1 nuclease (1U/ µl) with water to make the volume to 90µl. The digestion reaction was left at room temperature for 30 minutes and the enzyme deactivated by incubating at 70°C with 6 µl 0.5M EDTA. The DNA was then fragmented by nebulisation at 30psi for 180 seconds. The DNA was sequenced using the 454 Junior using the manufacturer’s recommended methods.

### Data analysis

Data analysis was performed in three stages, first host and contaminating reads were removed before abundance trimming, secondly reads were classified using lowest common ancestor (LCA) analysis and finally classified reads were assembled and analysed.

Initially reads were mapped using Newbler (Roche diagnostics) standard parameters against host and a local contamination library consisting of sequenced negative controls and reads identified as contamination in previous runs. Unmapped reads were written into a new fastq file using a custom python script and taken forward for further analysis. The remaining reads were abundance trimmed using Khmer^33^ using two passes to a maximum depth of 50, to remove over represented reads produce by the ɸ29 MDA method. Following this the reads were error trimmed using Prinseq^34^, with ends being trimmed to a Q20 cut off.

Blastn was then used to assign the reads against the NCBI nt database. Once completed lowest common ancestor (LCA) analysis was performed using MEGAN^35^, and reads associated with the bacterial species of interest were extracted to a new fastq file. These reads were then *de novo* assembled using standard parameters in SPAdes^36^ assembly outputs were assessed using QUAST (Quality Assessment Tool for Genome Assemblies)^34^. Reference assemblies were performed using the closest reference sequence identified using the LCA analysis. Antibiotic prediction was performed using Mykrobe^11^ for *S. aureus* and ResFinder^37^ for *E. Coli* using the *de novo* assembled reads.

## Results

### Sample processing

When the full blood processing method was applied to *E. coli* and *S. aureus*, good survival rates were found for both bacteria through-out all stages. Details of survival at each stage of processing can be found in Table 1

### Sequencing results S. aureus

After processing and DNA amplification the concentration of DNA was 139ng / µl, after sequencing the number of reads passing basic filter was 128,500. Once known contaminants were removed and error and abundance trimming complete 124,145 reads remained. After reads were assigned using Blastn and LCA analysis performed using MEGAN, 92.3% of reads were classified. 72.22% of reads were classified as Staphylococcus species and 62.1% reads were identified as *S. aureus*. 6.76% of the reads were identified as the genus Equus, and 1.72% were identified as *Parascaris equorum*. Remaining reads were not classified above the Class level in LCA analysis. When examining the *S. aureus* reads closer 4254 reads were identified to the subspecies level, (*Staphylococcus aureus subsp. aureus* HO 5096 0412), this subspecies has a complete genome available (GenBank GCA_000284535.1) and was used as a reference for reference mapping. Reference mapping covered 92% of the reference genome. When the reads identified as *Staphylococcus* using the LCA analysis were extracted and the reads *de novo* assembled 1212 contigs were produced. When this was compared to the same reference as the reference assembly 83% of the genome was covered with 10 misassemblies and a N50 of 3882. Mykrobe analysis was performed using the *de novo* assembly gave genotypic result for 12 antibiotics. When comparing the genotypic and phenotypic (BD Phoenix™) results matched in 11 of the 12 antibiotics (Table 2). The results for ciprofloxacin were inconclusive in genotypic tests, but resistant by phenotypic methods.

**Table 2.** phenotypic and genotypic result for antibiograms of *E. coli* (ResFinder) and *S. aureus* (Mykrobe)

|  | Phenotypic-(MIC) | Genotypic | Comment |
| --- | --- | --- | --- |
|  | *E. coli* |  |  |
| Ampicillin | R (>8) | R | *blaCMY-2* |
| Ceftriaxone | R (>4) | R | *blaCMY-2* |
| Ciprofloxacin | R (>1) | R | *gyrA* |
| Colistin | S (<=0.5) | S |  |
| Gentamicin | S (<=0.25) | S |  |
| Levofloxacin | R (>2) | R | *gyrA* |
| Meropenem | S (<=0.25) | S |  |
| Nalidixic acid | (>16) | R | *ermB* |
| Nitrofurantoin | S (<=16) | S |  |
| Trimethoprim | R (>4) | R | *dfrA* |
|  | *S. aureus* |  |  |
| Ciprofloxacin | R (>2) | Inconclusive |  |
| Clindamycin | S (<=0.25) | S |  |
| Erythromycin | S (<=0.25) | S |  |
| Fusidic acid | S (<=1) | S |  |
| Gentamicin | S (<1) | S |  |
| Methicillin | R (oxacillin 2) | R | *MecA* |
| Mupirocin | S (<=1) | S |  |
| Penicillin G | R (> 0.25) | R | *blaZ* |
| Rifampicin | S (<=0.25) | S |  |
| Tetracycline | S (<=0.5) | S |  |
| Trimethoprim | S (<=1) | S |  |
| Vancomycin | S (1) | S |  |

### Sequencing results E. coli

After processing and amplification, the DNA concentration of the sample was 143 ng/ µl. Post sequencing 173,597 reads passed the initial filter, once contaminants were removed and the reads error and abundance trimmed 170,243 reads remained. Overall 73% of all reads remaining after the analysis pipeline were identified as *E. coli*. 31959 (18%) reads had no identity. Unlike *S. aureus* it was not possible to type the *E. coli*, as 150 reads were assigned to O7:K1 and 211 reads were assigned to JJ1886. Reads which were identified as *Enterobacteriaceae, Escherichia* and *E. coli* by BLAST and LCA analysis were extracted from the fastq produced after pipeline completion. The genome of JJ1886 was available from the Integrated Microbial Genomes database (ID 2558309052), and the chromosomal sequence was used as the reference against which the extracted reads were assembled. After reference mapping assembly 93.5% of the genome was covered. When the reads were *de novo* assembled, 89% of the same reference was covered in 1334 contigs. The *de novo* assembly was used to identify several resistance markers using ResFinder. The resistance markers included *dfrA*, conferring trimethoprim resistance, *gyrA* conferring resistance to fluoroquinolones. *mdtK* is an efflux pump conferring resistance to norfloxacin and *blaCMY-2* which is an AmpC, conferring resistance to beta-lactams including cephalosporins. Additionally, eight drug efflux systems were identified. These results were concordant to the phenotypic data (Table 2).

## Discussion

Direct diagnostics using WGS from clinical samples would, in many ways, provide the ideal diagnostic method. By providing all the information from whole genome data with the speed of direct sample testing. There are increasing references to using metagenomic approaches to detect pathogens directly from clinical samples. A recent study, using the Oxford Nanopore Technologies MinION to sequence directly from TB smear positive respiratory samples, demonstrated it was possible to get phenotypically concordant results from sequencing in 12.5 hours^38^. Direct sequencing from orthopaedic device infections showed that a metagenomic sequencing approach gave species level specificity and sensitivity of 88% compared to traditional culture techniques^39^. In a study of ocular eye fluid samples, it was possible to match 9/12 bacteria using metagenomic sequencing compared to traditional culture techniques, with the additional advantage of being able to obtain sequence types and antibiograms for some of the identified pathogens^40^. Whilst these studies provide good evidence of the application of metagenomic sequencing, applying these methods directly to blood still has various are various obstacles, including low pathogen numbers, high host background and difficulty in interpreting genotypic data.

Here, a model was prepared to demonstrate the potential for direct from sample sequencing from whole blood. This method allows DNA to be ready within 3.5 hours of sample receipt, including bacterial isolation and DNA amplification. Once the DNA is prepared it could be processed on any sequencing platform. Final time to results would depend on the platform used, and the scale of batching required to make the method cost effective. A recent study looked at real time analysis of sequencing produced on the Oxford Nanopore MinION platform, showed it was possible to identify the bacterial species in a sample within 30 minutes of starting the run. Initial resistance predictions were ready within two hours, and all analysis was completed within 10 hours^41^. Combining good sample preparation and real-time analysis could allow results to be obtained in a clinically actionable time frame. In addition to the wet lab model, an analysis pipeline was developed to allow accurate interpretation of the results. This included removal of contamination, normalisation of reads and LCA analysis. Reference mapping and *de novo* assembly programmes were selected to give the best results for the data produced, in terms of amplification using ϕ29 MDA and sequencing on the 454 Junior. Fresh whole horse blood was used to model bacteraemia, as it was readily available. The process was developed to remove RBCs early in the process, as they represent the largest proportion of the cellular makeup of blood (up to 96%^42^), debulking the sample, and preventing release of oxidative agents. Selective lysis on WBCs allowed the release of any intracellular pathogens and exposed host nucleic acid to nucleases. *S. aureus* and *E. coli* were chosen due to different cell wall types and cell morphology; both demonstrated good survival during the developed sample processing.

The use of ϕ29 MDA was selected due to its ability to quickly and accurately amplify DNA without the need for specific primers, allowing its application to a wide range of pathogens. ϕ29 MDA has been associated with amplification bias, especially when starting from low input samples, previously this was thought to be random, but has now been shown to be systemic^43^. Lower reaction volumes have been shown to lower the effect of the bias, however significant scaling down from the current 5 l starting material would be technically difficult. Use of digital droplet MDA has been shown to give a more uniform coverage across the genome and lower none specific amplifcation^44^, however this requires specialised equipment. For this study amplification bias was compensated for bioinformatically, using a process of digital normalisation^33^, which removed over represented reads, and added an additional error removal stage by removing very low level k-mers. This aided in assembly of reads, as most assemblers assume even coverage across the genome, it also allowed the data size to be reduced without loss of information. Despite the bias in the amplification, good genome coverage and characterisation of the bacteria were achieved (89% and 93% gnome coverage using *de novo* methods), with missing sections often being repeat elements within the genome.

The majority of sequencing reads from spiked horse blood were associated with the spiked bacteria (62.1% *S. aureus* and 73% *E. coli*), considering the horse genome is over 500 times larger than the *E. coli* genome this shows that the vast majority of the host material was removed. Overall there was good concordance of phenotypic and genotypic results showing the potential for rapid genotypic prediction of antibiotic resistance from 10 bacterial cells in 1 ml of host blood. In the *S.aureus* sequencing the closest sub-species identified was *Staphylococcus aureus subsp. aureus* HO 5096 0412, this most likely represents the closest sequence in the reference database, rather than the actual type of this organism. Antibiotic resistance predictions were concordant with the phenotypic results except for ciprofloxacin. The inconclusive ciprofloxacin results demonstrate the need for improved understanding of the mechanisms of resistance. Ciprofloxacin resistance is harder to predict as it isn’t associated with a single gene acquisition. Three mechanisms of fluoroquinolone resistance have been proposed in *S. aureus*, Topoisomerase IV gene mutations, DNA gyrase gene mutations and an active efflux pump (NorA)^45^. The complexity of predicting ciprofloxacin resistance suggests that the database may be lacking in its ability to predict ciprofloxacin resistance, and so this is the most likely cause of the inconclusive result for ciprofloxacin resistance. The developers of Mykrobe, a commonly used antibiotic resistance predictor (Bradley et al^11^) found a false negativity rate of 4.6% for ciprofloxacin resistance.

In addition to identifying the isolates resistance to beta-lactams the database was able to identify the *blaZ* gene and *MecA* gene. Genotypic testing will never entirely replace phenotypic susceptibility testing, due to its inability to identify novel resistance determinants and the comprehensive nature of phenotypic testing. However, in the case of invasive sepsis, the gain in speed provided by not having to culture the organism to determine susceptibility could be life-saving.

Multiple antibiotic resistance factors were identified in the *E. coli*, which gave good concordance with the phenotypic data. Using the genotypic data, it was possible to rapidly identify the beta-lactamase present as *BlaCMY-2*. The rapid identification of the specific resistance genes in bacteria could help identify outbreaks by providing more information than a simple antibiogram. Additionally, it could help monitor resistance genes, to identify genes that are increasing in incidence. Large amounts of horizontal genome transfer amongst Gram negative bacteria has the potential to cause outbreaks of resistant bacteria through genes or plasmids^46^, which would be more complex to track. Rapid identification of genes causing the resistance in isolates could help inform epidemiological and outbreak studies which could involve several species of bacteria. A study of NGS sequencing data from bacteraemia isolates of *E. coli* have shown resistance prediction specificity of 97%^2^, if this was coupled with direct from sample sequencing genotypic prediction could inform treatment more rapidly than phenotypic test.

This study demonstrates that *de novo* assembly methods can be used to identify bacterial species and accurately predict antibiotic resistance. Lower coverage of the genomes after reference assembly (92% and 93%), could reflect that references used were close but not exact matches to the clinical isolates in the study, or the inability of short read assembly methods to assemble repeated elements which are longer than the read length. The inability to subtype the *E. coli* strain most likely reflects the large variety in *E. coli* genomes with the core genome representing only 13% of the pangenome^47^. An earlier study by the authors, using *E. coli* K12 MG1655 gave genome coverage with 100% sequence identity with the E. coli K12 MG1655 reference (data not shown). Clinical isolates rather than fully characterised laboratory strains were selected to test the applicability of the analysis to unknown samples. The ability to identify bacteria in a sample will be very dependent on the quality of the database used, and results will be biased towards common pathogens, but this method should be applicable to most bacteria found in clinical samples.

Limitations of this model include differences between horse and human blood, and difference in sample between spiked bacteria and bacteria causing a true infection. Additionally, factors in the blood such as immune and inflammatory reactions could alter the efficiency of the preparation method. As the bacteria was only quantified using dilution methods it is not possible to know the exact number of bacteria spiked into the sample. Whilst three replicates of each bacteria were checked for survival through the model, it was only possible to sequence one representative of each of the bacteria. However, this model shows proof of principle that direct unbiased sequencing directly from blood is possible and could be used to diagnose and inform treatment of bacterial bloodstream infections.

## Acknowledgements

This project was funded by Roche Diagnostics as a scientific studentship award and completed at Public Health England Centre for Infection.

Declarations of interest.

None

